# hudson: A User-Friendly R Package to Extend Manhattan Plots

**DOI:** 10.1101/2022.01.25.474274

**Authors:** Anastasia Lucas, Anurag Verma, Marylyn D. Ritchie

## Abstract

The Manhattan plot is one of the most widely used visualization techniques when plotting summary statistics from genome-wide or phenome-wide association studies. While there are a number of existing tools to create these plots, there is room for extending their utility to satisfy increasingly complex and comprehensive analyses as well as the need for comparisons between different sets of results or between discovery and replication datasets. The R package presented here, *hudson*, provides user-friendly plotting functions intended for use with genome, phenome, and exposome-wide association studies, but its flexible framework can be utilized for a wide variety genome-wide analyses. Further, we have extended these figures to allow for interactive elements to facilitate results exploration for the ever-increasing large-scale dimensionality of these data. *hudson* can be obtained from https://github.com/RitchieLab/hudson.

## 1 Introduction

Manhattan plots displaying p-values at a genome scale have long been considered the standard method to provide a big-picture overview of genome-wide association study (GWAS) results. As genome-^1^, phenome-^2^, and exposome^3^ wide analyses (GWAS, PheWAS, EWAS) become more sophisticated and involve more dimensions of data, there is room to further develop and improve on data visualization techniques for the presentation of results. Including post-GWAS analysis methods and study designs that extend beyond basic GWAS, such as comparisons with previously published summary statistics, sex-stratified analyses, gene-based studies, transcriptome-wide association studies, etc., is becoming the norm in these large-scale scientific studies of the genetic or genomic architecture of complex traits. To perform comparisons of different datasets or different sets of results, using popular Manhattan plot packages such as qqman^4^ or PheWAS^5^ would require separate or side-by-side figures presented for comparison; these can be difficult to mentally map the chromosome positions, a task necessary to notice key similarities and differences in loci. It can also be challenging to drill down to specific results as most of these packages produce static images that make it difficult to discern specific data points.

While there exist several packages such as miamiplot^6^, gwaRs^7^, and karyoploteR^8^, for comparing two Manhattan plots using SNP data from association studies, many are inflexible to data that deviates from a traditional GWAS study and are constrained to a static figure that is limited to three or four attributes—x-axis position, y-axis position, color, and/or shape. To the authors’ knowledge, there are no maintained Manhattan plot packages that include methods for creating mirrored Manhattan plot figures as well as allow for data that cannot be organized into chromosomes, such as exposures or tissues, and include methods to create interactive figures that allow the user to view meta data without the need for heavy coding. Here, we present *hudson*, an R package to create mirrored Manhattan plots for various data types with added capabilities to create interactive figures with the goal of providing a single easy to use data visualization resource for researchers.

## 2 Methods

*hudson* was developed as an open-source R package. The plotting mechanisms are dependent on the widely used graphing library ggplot2^9^ and the interactive functions utilize the interactive ggplot extension, ggiraph^10^. Additionally, *hudson* will make use of ggrepel^11^, an R library that automatically repels data labels to prevent overlapping text, if it is installed on the user’s system. The package consists of seven main functions, two each designed for both static and interactive visualization of GWAS, PheWAS, and EWAS analysis comparisons and also a QQ-plot function. Sometimes referred to as Miami plots^12,13^ or Chicago plots^14^ our Hudson plots are characterized by a divergent y-axis and a shared x-axis, allowing for a position by position, or more generally, variable by variable, comparison of data.

In their most basic form, each half of the Hudson plot consists of a typical Manhattan or PheWAS Manhattan plot. The x-axis generally represents either 1) chromosome and base pair location across the genome or 2) phenotype/disease categories. From there, users can choose to set multiple parameters to highlight and/or annotate above certain p-value thresholds or by name, add threshold lines, and make various theme changes such as changing the background or colors, turning on or off x-axis marker blocks, and manually adjusting the y-axes. The functions designed to work with genomic data can also accept any number of chromosomes such that the user can plot non-human genomes or any subset of interest such as specific gene sets as shown in Drivas et al 2021^15^.

Users wishing to make interactive versions of figures provide custom annotations in additional metadata columns. This metadata will be displayed when the user hovers over a point on the figure. This allows a user to interact with the data and for each point determine the characteristics of that data point (for example, SNP ID, gene ID, odds ratio or beta, and p-value). Further, the user can provide a link that should be opened in a browser window when the user clicks on the point; this could link out to the dbSNP information about that variant, OMIM information about that gene, or a publication in PubMed for example. These features eliminate the need to perform manual lookups when investigating SNPs or genes of interest. The interactive figures can then be saved as HTML files which can easily be shared among researchers and opened locally in any internet browser.

As one of the goals of the package is to produce publication quality images, we have considered the need for color-blind friendly color palettes. By default, *hudson* will select colors from a 15-color palette that is distinguishable for people with deuteranopia and protanopia, and generally safe for tritanopia^*^. If more than 15 colors are required, *hudson* will interpolate colors from Google AI’s Turbo palette^†^ which is both perceptually uniform and distinguishable for people with all forms of partial colorblindness, offering a benefit over a traditional rainbow palette. Although users will be able to override these palettes and specify their own color scheme, we hope that having them built in as defaults will make it easier for users to create more widely accessible figures.

Though we have provided a flexible framework for visualizing many types of data, we recognize that there will be instances where the user requires further customization or features and, therefore, have made the source code for both packages freely available on GitHub under the GPL-3 license.

Additionally, for the most part, each function is able to be downloaded, modified if needed, and run as a standalone script in R. For example, in Veturi et al. 2021, we modified *hudson* code to generate Figure 3 in which we rotated the plot 90 degrees so that we could display two Hudson plots side by side to compare gene expression based PheWAS (Xpress-PheWAS) and SNP-based PheWAS in two datasets^16^. The package was designed this way intentionally to increase accessibility to researchers who wish to make small tweaks to their figures, but do not have as much experience with R package development.

## 3 Usage

Figure 1 provides a more comprehensive example of the a few of the use cases for the *hudson* package using summary statistics obtained from the NHGRI-EBI GWAS Catalog^17^ and the default data from the *hudson* package. Panel A displays standard Hudson plot with GWAS results from the Ehret et al. blood pressure GWAS^18^ for two traits, systolic blood pressure (SBP) and diastolic blood pressure (DBP). Panel B displays all four phenotypes from the Spracklen et al. lipid GWAS results^19^ along with the betas for X in the top and Y in the bottom. Panel C displays an example of an exposome-wide association study using a dummy dataset provided by the *hudson* package. Panel D displays a gene-based PheWAS plot. These examples were selected to demonstrate the wide variability of types of data that can be displayed in Hudson plots.

**Figure 1:**
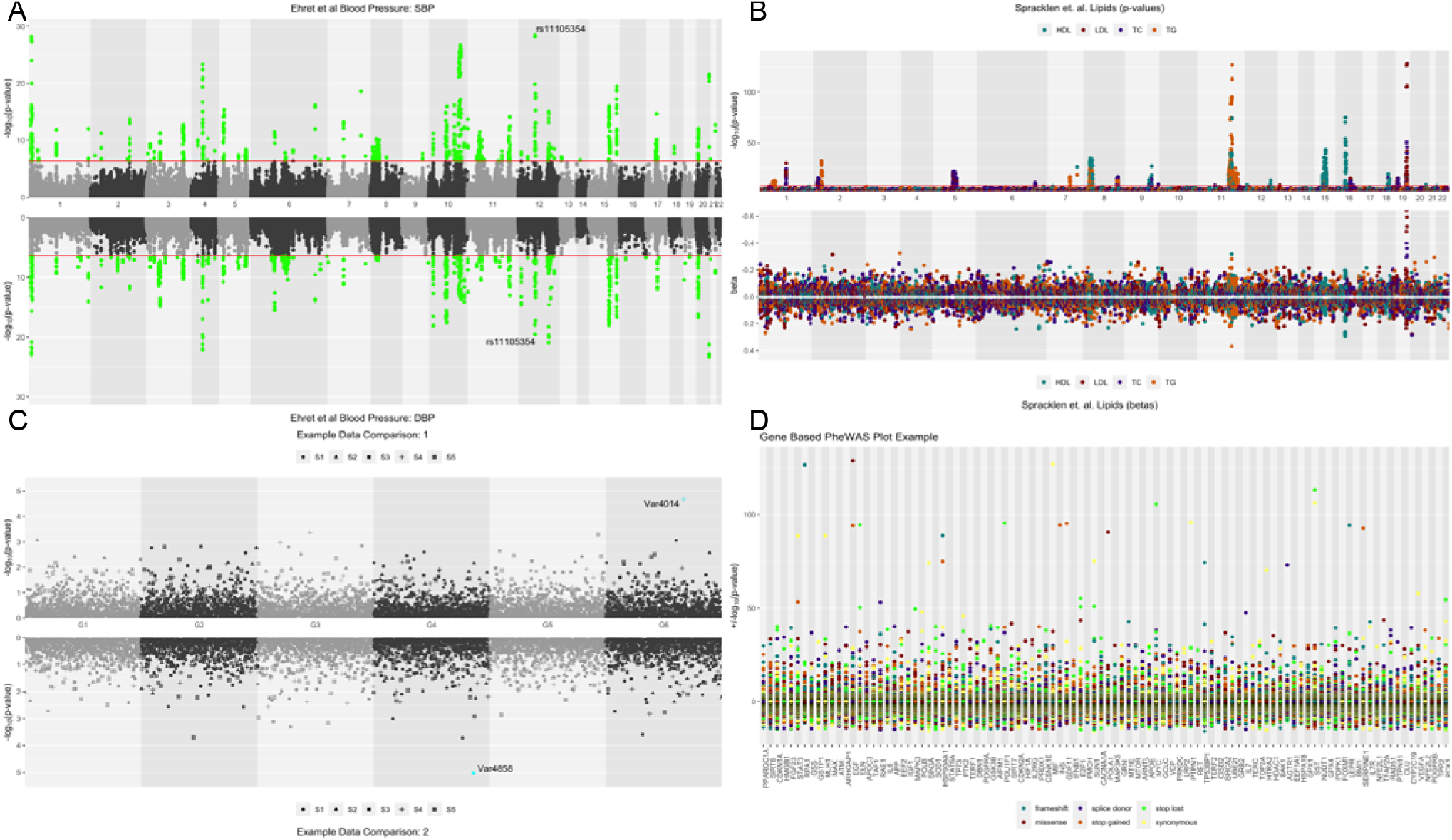
Examples of GWAS (A), PheWAS (B), and EWAS (C) hudson plots, each showing different use cases for the *hudson* package. Panel D shows an example of a modified version of the PheWAS plotting method that organizes variants by gene in lieu of chromosomes. Note the complete lack of linkage disequilibrium characteristic of fabricated data in comparison with Figures 2–5 of Drivas, et al.

Figure 2 shows an example of an interactive PheWAS Hudson plot figure. This plot includes the data from the Spracklen et al paper. The methods used to create the individual panels for each of these figures are explained in more detail below and the code used to create the figures can be found at https://github.com/RitchieLab/utility/tree/master/personal/ana/hudson-paper.

**Figure 2:**
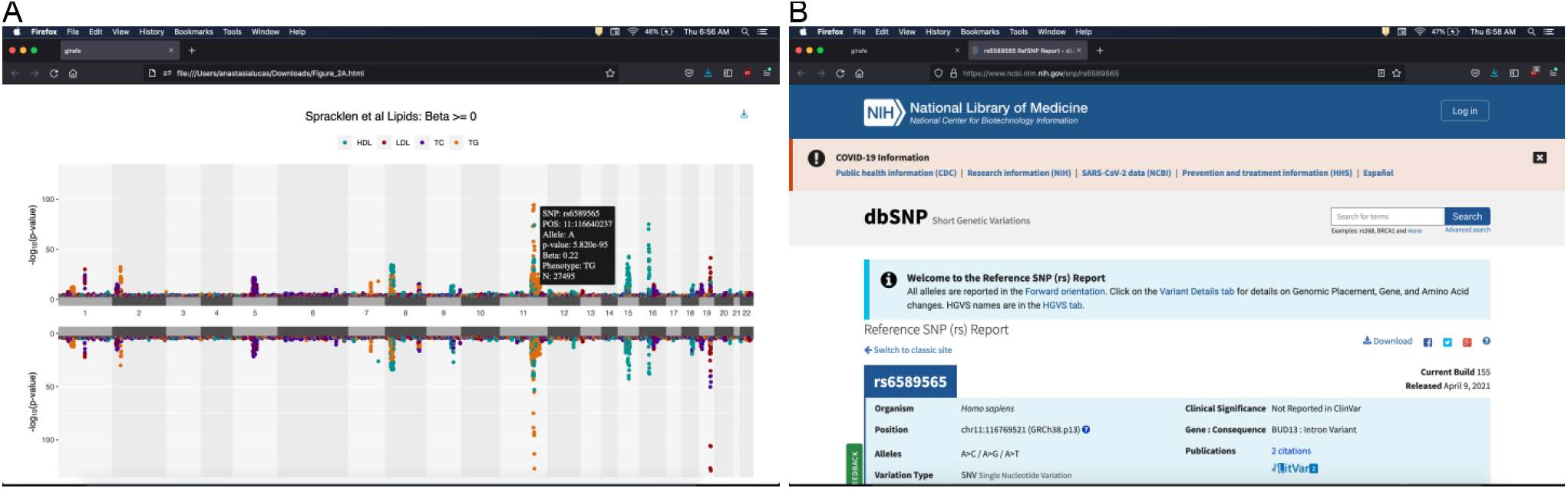
Screenshot of an interactive PheWAS Hudson plot being viewed in an internet browser. Panel A shows an example of annotations that will appear when a point corresponding to a SNP is hovered over while panel B shows the dbSNP entry for that variant opening in a new browser window when the user clicks on the point.

### 3.1 Mirrored GWAS plots

GWAS Hudson plots produced by the *gmirror()* function will accept as input genomic data such as those output from common GWAS analysis tools like PLINK^20^. Thus, data should contain at least columns for variant name, chromosome, position, and p-value (or other numeric value, such as beta). As in the other plotting functions, an additional layer of information can optionally be included in column mapped to shape. Figure 1A contains an example of a GWAS Hudson plot using summary statistics from the GWAS Catalog for studies GCST006258^18^ and GCST006259^18^ downloaded on 11/22/2021. This example makes use of *hudson’s* p-value threshold highlighting and variant annotation options.

### 3.2 Mirrored PheWAS plots

PheWAS Hudson plots are produced by the *phemirror()* function and take input genomic data similar to described above, just with the addition of a phenotype column that will be automatically mapped to a color. Note that this “phenotype” could be any attribute the user wishes to map to color, for example variant annotations, such that this is not solely limited to visualizing data from a PheWAS analysis. Figure 1B shows an example of visualizing two different data types from a collection of four traits from a lipid association study^19^ (GCST004232, GCST004233, GCST004235 and GCST004237) downloaded from the GWAS Catalog on 10/24/2021. Here we see p-values in the upper panel and beta in the lower panel, making use of *hudson*’s free y-axis feature. This panel also shows the use of chromosome panels and base blocks to better distinguish which points belong to which chromosome.

### 3.3 Non-genomic and other plot types

Although we have referred to the types of figures the *emirror()* function can create as EWAS plots, this function is suitable for any analysis where many results are organized into categories, such as tissues or pathways, provided there is a common x-axis between the upper and lower panels. Thus, only three columns of information are needed: variant, group, and p-value or other numeric value. Figure 1C is an example of a plot using the simulated EWAS data provided by the package with an additional attribute mapped to shape. Here, we again can use similar variable (and/or p-value) highlighting thresholds and theme options as described above.

To highlight the flexibility of the package, we have also included an example that is not made by the default package, but was obtained with only minor modifications to the source code in Figure 1D. This version of the mirrored PheWAS plot uses gene names as our pseudo ‘chromosome’ grouping, achieved with the addition of a single parameter to specify ordering, and was further modified to use a fixed position for the variants rather than a number line. The code used to create this figure can be found at https://github.com/RitchieLab/utility/blob/master/personal/ana/scripts/phegene_hudson.R.

### 3.4 Interactive plots

Users wishing to make interactive versions of the figures can provide data in the same input format described above to analogous functions prefixed with ‘i’, i.e. *igmirror()*, *iphemirror()*, and *iemirror()*. However, here they may optionally provide custom annotations in an additional metadata column.

This metadata will be displayed when the user hovers over a point on the figure as shown in the screenshot of an interactive PheWAS Hudson plot being viewed in an internet browser in Figure 2A. Various user-defined annotations including the variant name, chromosomal position, beta, effect allele, and number of samples can be added. Further, the user can provide a link that should be opened in a new browser window when the user clicks on the point, as demonstrated in Figure 2B. This link could be anything from a dbSNP query to a Google search. Both the annotations and connections to external databases eliminate, or at least greatly reduce, the need to perform manual lookups when initially investigating SNPs, genes, or other variables of interest. The interactive figures are then saved as HTML files which can easily be shared among researchers and can be opened in any internet browser.

## 4 Discussion

The *hudson* package allows users to easily create publication quality or shareable interactive mirrored Manhattan plot figures. When performing association analyses on hundreds of thousands or even millions of variables, it can be a challenge to interpret the results from large tables alone. Visualizing the results in different ways and specifically with the ability to compare different groups in a single plot with accurate alignment of the variables can make this interpretation process significantly more accurate and efficient. In *hudson*, we provide a flexible, open-source framework that allows users to visualize data from a wide variety of study designs as well as annotate those figures with interactive metadata, while also encouraging more advanced users to customize the scripts to meet individual research needs that are not explicitly fulfilled by the base package.

## Author Contributions

AL conceived of and developed the software package and wrote the manuscript. AV made design contributions to the package. MDR supervised the project. All authors contributed to editing the article and approved the submitted version.

## Acknowledgments

We acknowledge GitHub user @dbaranger who published R code for the Turbo color palette implemented in the package.

## Data Availability Statement

Example datasets used in this study can be found in the GWAS Catalog (https://www.ebi.ac.uk/gwas/ and toy data can be found at Ritchie Lab GitHub (https://github.com/RitchieLab/hudson/tree/master/data) or loaded from the package itself.

## Footnotes

* Kryzwinski, 2012 http://mkweb.bcgsc.ca/biovis2012/

† Mikhailov, 2019 https://ai.googleblog.com/2019/08/turbo-improved-rainbow-colormap-for.html

